# Resting-state Co-activation Patterns as Accurate Predictors of Alzheimer’s Disease in Aged Mice

**DOI:** 10.1101/2020.10.22.349852

**Authors:** Mohit H. Adhikari, Michaël E. Belloy, Annemie Van der Linden, Georgios A. Keliris, Marleen Verhoye

**Affiliations:** Bio-imaging Lab, Department of Bio-medical Sciences, University of Antwerp, Antwerp, Belgium

**Keywords:** Alzheimer’s disease_1_, resting-state fMRI_2_, co-activation patterns_3_, predictive modeling_4_, classification_5_

## Abstract

Alzheimer’s disease (AD), a neurodegenerative disorder marked by accumulation of extracellular amyloid-beta (Aβ) plaques leads to progressive loss of memory and cognitive function. Resting state fMRI (RS-fMRI) studies have provided links between these two observations in terms of disruption of default mode and task positive resting state networks (RSNs). Important insights underlying these disruptions were recently obtained by investigating dynamic fluctuations in RS-fMRI signals in old TG2576 mice (mouse model of amyloidosis) using a set of quasi-periodic patterns (QPP). QPPs represent repeating spatiotemporal patterns of neural activity of predefined temporal length. In this article, we used an alternative methodology of co-activation patterns (CAPs) that represent instantaneous and transient brain configurations that are likely contributors to the emergence of commonly observed resting state networks (RSNs) and QPPs. We followed a recently published approach for obtaining CAPs that divided all time frames, instead of those corresponding to supra-threshold activations of a seed region as done traditionally, to extract CAPs from RS-fMRI recordings in 10 TG2576 female mice and 8 wild type littermates at 18 months of age. Subsequently we matched the CAPs from the two groups using the Hungarian method and compared the temporal (duration, occurrence rate) and the spatial (lateralization of significantly activated voxels) properties of matched CAPs. We found robust differences in the spatial components of matched CAPs. Finally, we used supervised learning to train a classifier using either the temporal or the spatial component of CAPs to distinguish the transgenic mice from the WT. We found that while duration and occurrence rates of all CAPs performed the classification with significantly higher accuracy than the chance-level, blood oxygen level dependent (BOLD) signals of significantly activated voxels from individual CAPs turned out to be a significantly better predictive feature demonstrating a near perfect classification accuracy. Our results demonstrate resting-state co-activation patterns are a promising candidate for a diagnostic, and potentially, prognostic biomarker of Alzheimer’s disease.

## 1 Introduction

Alzheimer’s disease (AD) is a neurodegenerative disorder that causes progressive loss of learning abilities, memory, and overall cognitive function. The characteristic features of the disease are the accumulation of extracellular amyloid-beta (Aβ) plaques and intracellular neurofibrillary tangles. In order to understand how the accumulation of plaques could contribute to the development of AD symptoms, it is important to investigate changes in neural activities especially at the network or whole-brain level. Resting-state functional magnetic resonance imaging (RS-fMRI) has been instrumental in uncovering such global network-level changes across the whole-brain in several neurological and neuropsychiatric disorders such as stroke (Baldassarre et al. 2014; Siegel et al. 2016; Carter et al. 2010), coma (Di Perri et al. 2018; Chennu et al. 2017), depression (Drysdale et al. 2017). In case of AD, the disruption of the default mode and task-positive networks have been identified as promising markers of the disease. Specifically, alterations in the default-mode network (DMN) functional connectivity (FC) have been correlated with increases in amyloid-beta levels (Greicius et al. 2004; Li and Wahlund 2011).

Traditional analyses of RS-fMRI signals have involved calculation of static (seed-based or pairwise between regions of interest) FC. FC estimates correlations of BOLD signals of regions from the entire scanning period disregarding the variations in FC during the scan. However, several recent studies (Hutchison et al. 2013; Deco and Kringelbach 2017; Hindriks et al. 2016) have shown that temporal fluctuations in FC within the scan can inform on interplay between various brain states. Several methods have been proposed to extract this dynamic information in the resting-state FC. The most-straightforward one uses a sliding window approach. Here, whole-brain FC is calculated in a time window of fixed duration that is then moved over the entire scan in order to obtain a series of FC values over the whole scan (Hutchison et al. 2013). Test-statistics are then calculated using this FC time series and compared against the null hypothesis of stationarity (Hindriks et al. 2016). Another approach consists of a point-process analysis (Liu and Duyn 2013; Liu et al. 2018) in which fMRI time frames where signal of a given region of interest (i.e. seed) crosses a specific percentile threshold, are clustered to identify different co-activation patterns (CAPs). Voxel-wise activation pattern, averaged across these selected frames (typically only 15% of the total) matches very closely with the seed-based correlation maps obtained using all frames. CAPs represent transient brain states that are believed to contribute to the emergence of resting-state networks (RSNs) found in the static FC estimation (Liu et al. 2018). Recently, Gutierrez-Barragan et al. used a modified approach in which they clustered all time frames from RS-fMRI scans in mice and found six robust CAPs in different datasets (Gutierrez-Barragan et al. 2019).

In this paper, we used the methodology of Gutierrez-Barragan et al., to identify CAPs in a cohort of old (18-months) TG2576 (mouse-model of amyloidosis) mice and their age-matched control. In this cohort, Belloy et al. identified changes in a set of recurring spatio-temporal patterns of neural activity of predefined temporal length called the quasi-periodic patterns (QPPs) (Belloy et al. 2018). We compared the spatial and temporal components of CAPs between the two groups. Subsequently, we hypothesized that the CAP properties will accurately distinguish the transgenic animals from healthy controls and argue that it could be effective in the development of a biomarker for Alzheimer’s disease.

## 2 Materials and Methods

All the data analyzed in this manuscript were originally acquired and published in an earlier manuscript (Belloy et al. 2018). The acquisition and processing steps are included here for completeness.

### Ethical statement

All procedures were performed in strict accordance with the European Directive 2010/63/EU on the protection of animals used for scientific purposes. The protocols were approved by the Committee on Animal Care and Use at the University of Antwerp, Belgium (permit number 2014-04) and all efforts were made to minimize animal suffering.

### Animals

The TG2576 mouse model of amyloidosis overexpresses the human mutant form of amyloid precursor protein (APP), which carries the Swedish mutation (KM670/671NL), controlled by the hamster prion protein promoter (Hsiao et al. 1996). Aβ plaque development starts at the age of 9–11 months (Hsiao et al. 1996), while plaque burden increases markedly with age (Kuo et al. 2000). The cohort used in this study consisted of 10 female TG2576 (henceforth referred to as TG) mice at the age of 18 months and 8 age-matched wild-type (WT) littermates. RS-fMRI data were collected while the animals were under an anaesthesia protocol comprising 0.4 % isoflurane, a bolus injection of medetomidine (0.3 mg/kg), and a subcutaneous infusion of medetomidine (0.6 mg/kg/h).

### MRI procedures and functional scan pre-processing

MRI scans were acquired on a 9.4 T Biospec system, with a four-element receive-only phase array coil and a volume resonator for transmission. Structural images were acquired in three orthogonal directions, using Turbo Rapid Imaging with Refocused echoes (RARE), for reproducible slice positioning (repetition time 3000 ms, effective echo time 33 ms, 16 slices of 0.4 mm). B0 field maps were acquired, followed by local shimming. RS-fMRI scans were acquired with a gradient-echo echo-planar imaging (EPI) sequence (field of view (20×20) mm^2^, matrix dimensions [128×64], three slices of 0.4 mm, flip angle 55°, bandwidth 400 kHz, repetition time 500 ms, echo time 16 ms, 2400 repetitions). High temporal resolution was required to investigate temporal fluctuations in the data. Due to resultant technical limitation, slice number was restricted to three. Slices were positioned 0.1 mm caudally of bregma, according to the Paxinos and Franklink stereotaxic mouse brain atlas (Franklin and Paxinos 2013).

Motion parameters for each functional scan were obtained using 6 rigid body parameters. Images were realigned and normalized to a user-defined reference subject, followed by smoothing (σ = 2 pixels). During image normalization, intensities of outer slices are partially lost. Analyses were thus restricted to the single center slice (MATLAB2017b). Motion vectors were then regressed out of the image-series. These procedures were performed using Statistical Parametric Mapping (SPM12) software (Wellcome Department of Cognitive Neurology, London, UK). Images were then filtered using a 0.01-0.2Hz FIR band-pass filter, quadratic detrended and normalized to unit variance. Transient time points at the start and end of the image-series were removed before and after filtering. For the detection of CAPs, a brain mask was employed to exclude the contribution of the ventricles. Global signal regression was not carried out.

### Extraction of CAPs

As mentioned in the introduction, we followed the approach by Gutierrez-Barragan et al. 2019 to obtain the co-activation patterns in each group (WT and TG). Thus, we first concatenated the filtered images from each animal in the group to form a group-level image-series. We then clustered all time frames in this image-series using K-means++ algorithm by assessing their spatial dissimilarity with each other in terms of correlation distance (1 - Pearson’s correlation). Clustering was done for a range of clusters between 2 and 20 and in each case, we calculated across-subject variance explained by the clustering solution as follows (Goutte et al. 1999):

1. Between cluster variance, 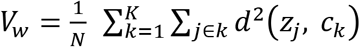 where, *N* is the total number of observations (time frames); *K* is number of clusters and *d* denotes the Euclidean distance between the global centroid *c_k_* of the *k^th^* cluster and *j_th_* observation belonging to the *k^th^* cluster.
2. Between cluster variance, 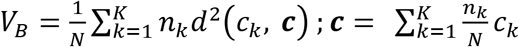 where *d* is the distance between the global centroid *c* and the cluster centroid *c_k_* and *n_k_* is the number of observations (time frames) in the *k^th^* cluster.
3. 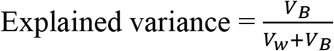

We then plotted the explained variance as a function of partitions of the image-series with increasing number of clusters (in the range from 2 to 20) and identified the minimum number at which the variance reached a saturation level (elbow point) as the optimal number of clusters. We confirmed the elbow point by making sure that the fractional gain in the explained variance for this partition with *k* clusters when compared to the partition with *k-1* clusters was less than 10% (Gutierrez-Barragan et al. 2019). Voxel-wise BOLD signal intensities were averaged across all time frames within each cluster to produce group-level CAPs. One-sample T maps corrected for multiple comparisons using the Bonferroni correction (p<0.01) were subsequently obtained for each CAP. We then calculated temporal and spatial properties of CAPs for each subject within each group:

1. Occurrence fraction: the ratio, for each CAP, of the number of time frames labelled with its id to the total number of frames within a subject. This is a subject-level measure.
2. Duration: number of consecutive frames corresponding to a CAP, averaged across all occurrences of the CAP within a subject; also a subject-level measure.
3. Laterality index: Difference in the average value of T-statistic per voxel, between left and right hemispheres, normalized by the average T-statistic per voxel in the entire brain. Here the T-statistic for each voxel is obtained by comparing its mean BOLD signal intensity, across all time frames within each subject belonging to the CAP, with zero using a one-sample T-test. Only voxels that whose activations (mean BOLD signal intensities) are significantly different from zero (p < 0.01; one sample T-test, Bonferroni corrected) were considered. The laterality index varies between −1 (completely right lateralized pattern) and 1 (completely left lateralized pattern) with 0 indicating a bilateral pattern with no preference for any hemisphere. We calculated two separate values of laterality for co-activations (T > 0) and co-deactivations (T < 0) respectively.

CAPs extracted from the image-series of both groups were spatially matched using the Hungarian method (Kuhn 1955) with 1-Pearson’s correlation taken as the distance metric. The strength of spatial similarity between every pair of matched CAPs was compared against a null hypothesis that it arises by chance. Thus, we shuffled randomly the CAP labels of all frames (thereby preserving the cluster size) from each group’s image-series and then obtained random surrogate CAPs by averaging across frames with the same CAP label. We then calculated the Pearson’s correlation between surrogate CAPs from each group while maintaining the matching found in the original datasets. We repeated this procedure 10000 times and built a surrogate distribution of correlation values and identified a threshold correlation value with p = 10E-5. All matched CAP pairs with canonical correlations falling below this value were not considered for further analyses/comparisons.

### Statistical comparisons

We compared for every pair of matched CAPs the medians, across subjects, of the properties mentioned above using a two sample rank-sum test, corrected for the number of comparisons using the Benjamini – Hochberg correction (Benjamini and Hochberg 1995) for controlling the false discovery rate (FDR). At first, we made these comparisons for every pair of matched CAPs (with Pearson’s correlation higher than the threshold from a null distribution of correlation values arising by chance) extracted from a specific partition of image-series with a fixed number of clusters. Subsequently, we compared the matched CAPs from all partitions with the number of clusters ranging between 2 and 20.

### Extraction of CAPs from a combined image-series

We also performed the CAP analysis on a combined image-series formed from a concatenation of both group-level image-series in order to avoid the necessity of matching. We compared the mean temporal and spatial properties between the groups for each combined CAP.

### Classification

Temporal and spatial components of CAPs were used as features in a supervised learning approach to distinguish TG animals from WT. We used a multinomial logistic regression (MLR) as a classifier and trained it on CAP features from 80% of the subjects and tested its accuracy on the remaining 20%. The regularization parameter in the MLR classifier was set to 10 to control for over-fitting. We repeated the accuracy calculation on 100 trials of randomly sampled train and validation sets and compared the mean accuracy with chance-level accuracy, averaged across 100 surrogate trials in each of which the identities of the subjects were shuffled while maintaining the size of each group (8 WT and 10 TG animals). Mean chance-level accuracy was expected to be ~ 45% which is the ratio of the size of the smallest class (WT) to the total number of subjects. We also computed the confusion matrices that give the true and false positive rates for each class and hence inform about sensitivity and specificity of the classifier. The confusion matrices were obtained using the true and predicted labels pooled from validation sets of all 100 trials.

We used the features of only those CAPs that showed a Pearson’s correlation of 0.5 and above between the two groups for training the MLR classifier. From all such CAPs within every partition, we pooled their (a) duration and occurrence rate, and, (b) BOLD intensities of voxels whose activations were found to be significantly different from zero in either the WT or the TG CAP. Each feature was z-scored across subjects so that their relative rankings were used for classification.

As CAPs were obtained using all subjects, information on validation set subjects could bias the classifier to predict them more accurately than otherwise possible. In order to avoid this bias, we extracted the group-level WT and TG CAPs only from the training set and then spatially correlated them with the image-series for every subject. Local peaks of the correlation time series were identified and voxel-wise averaged to use as initial centroids for the K-means clustering of all frames in the scan. The frames belonging to the same cluster were voxel-wise averaged to construct either a WT-like CAP or a TG-like CAP for every subject. Thus for each subject we obtained, for every group-level WT and TG CAP, two sets of features belonging to a WT-like CAP and a TG-like CAP. This was necessary to mimic actual situations in which the identity of a new test subject would not be known. We then trained the classifier using the WT-like and TG-like spatial and temporal features of the training set and tested its accuracy on the validation set. The whole-procedure was repeated for 50 trials and comparison of mean accuracy, across trials, with chance-level accuracy and calculation of confusion matrices were done as described in the paragraph above.

## 3 Results

### 3.1 Identification of Group-level CAPs

We began by partitioning the concatenated image-series of each group with the number of clusters ranging between 2 and 20. Following Gutierrez-Barragan et al. 2019, we calculated the across-subject variance explained by each partition and calculated the elbow point. As Figure 1A shows, the elbow point turned out to be a partition with 7 clusters as this was the first partition at which the fractional gain in explained variance fell below 10% for both groups (Figure 1B). We then took voxel-wise averages of BOLD signal intensities across all time frames belonging to each of the 7 clusters to obtain the group-level WT and TG CAPs. Figure 2 shows the T-statistic values for significantly (p < 1E-5, one-sample T-test, Bonferroni corrected) co-activated (T > 0) and co-deactivated (T < 0) voxels for each of the 7 CAPs matched using the Hungarian method between groups. CAPs were ordered in the descending order of spatial correlation between WT and TG groups.

**Figure 1:**
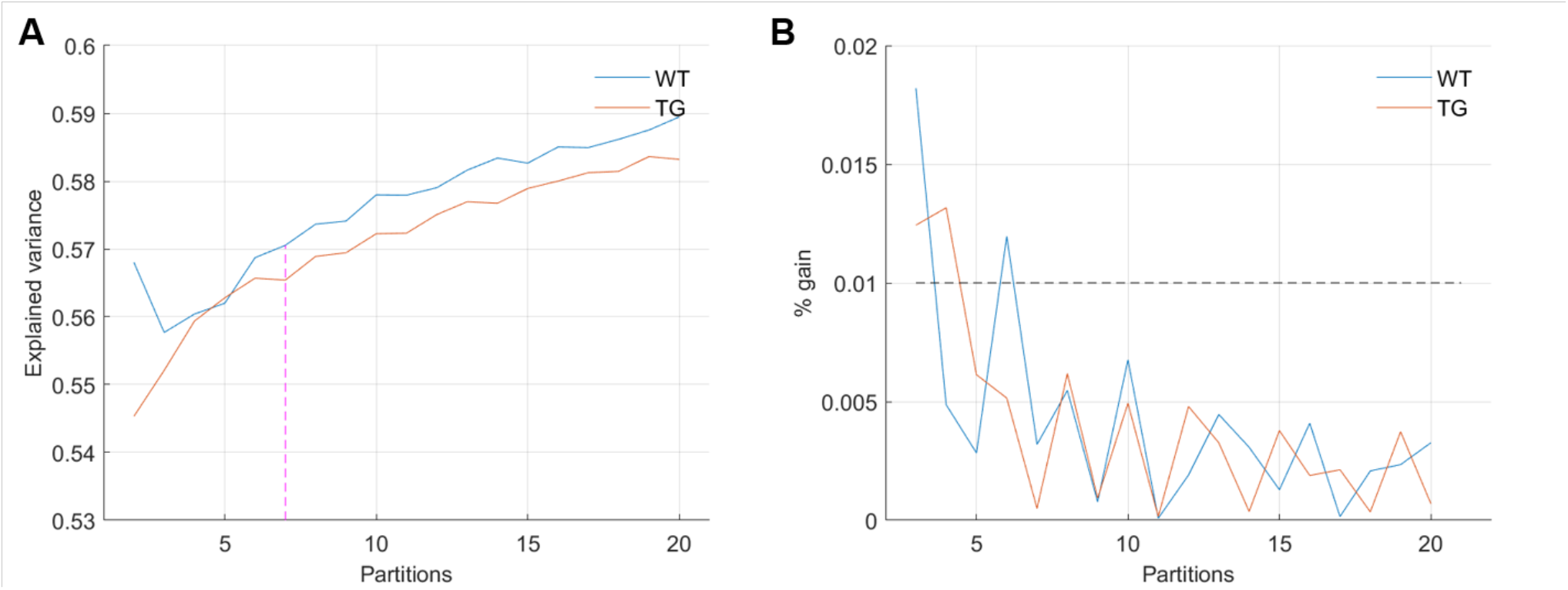
**(A)** Across-subject variance explained as a function of number of clusters in the partition of group-level image-series without global signal regression. The elbow point here beyond which the explained variance saturates is found to be at the partition with 7 clusters. **(B)** Fraction of gain in the explained variance as the number of clusters in the partition increase from k to k+1, as a function of partitions. The elbow point of 7 clusters is the first instance for both groups at which the fractional gain in explained variance falls below 10%.

**Figure 2:**
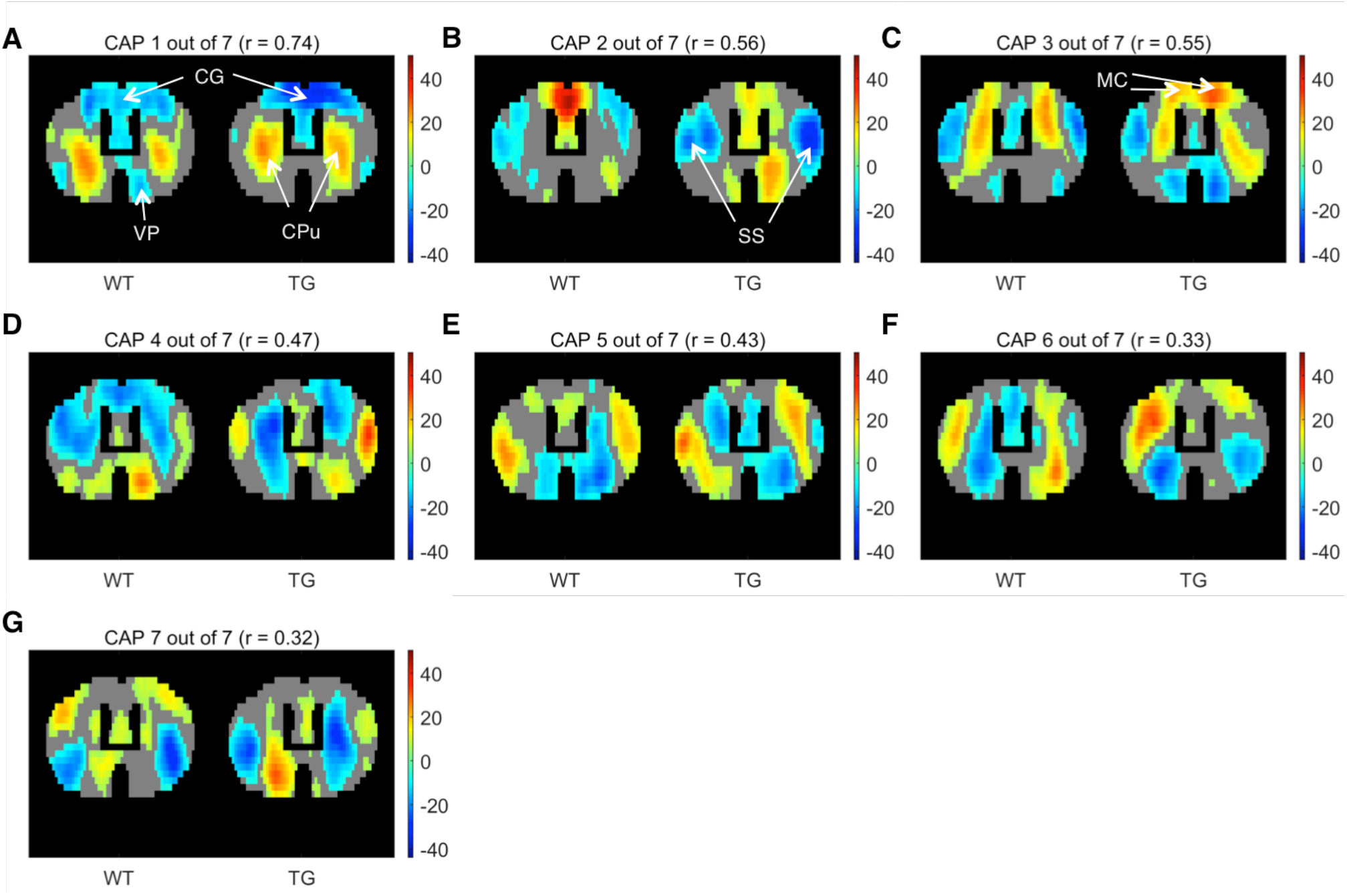
One-sample T-test maps showing significantly (p < 1E-5; Bonferroni corrected) co-activated (T > 0) and co-deactivated (T < 0) voxels for 7 WT CAPs and their corresponding matched patterns in the TG. CAPs are ordered in the descending order of spatial similarity between matched CAPs expressed in terms of Pearson’s correlation mentioned in the title of each pattern. CG: Cingulate cortex, CPu: Caudate Putamen, VP: Ventral Pallidum, SS: Somatosensory cortex, MC: Motor cortex.

CAP 1 was characterized by co-deactivations of cingulate (CG) and motor (MT) cortices and activations of dorsal and ventral caudate putamen (Cpu) of striatum. CAP 2 on the other hand was characterized by co-activations of mainly the CG and MT cortices along with co-deactivations of somatosensory (SS) cortices. Pearson’s correlation between CAP 1 and 2 was −0.36. CAPs 3 and 4 were similarly an anti-correlated pair of patterns (r = −0.4) characterized by simultaneous co-activations and co-deactivations, respectively, of CG, MT as well as the CPu that were anti-correlated with activations of somatosensory cortex. CAPs 5-7 displayed less commonly observed physiological configurations. Thus these CAPs featured characteristic regions – the CG and SS, respectively, that typically constitute the mouse DMN-like and latero-cortical RSNs (Liska et al. 2015; Gozzi and Schwarz 2016) respectively.

### 3.2 Comparisons of properties of matched CAPs

One-sample T-maps of CAPs in Figure 2 showed that while the matched CAPs had high spatial similarity, the co-activations and co deactivations were not necessarily symmetrical across hemispheres. Therefore, we used lateralization of co-activation and co-deactivation of voxels as a quantifiable metric to assess spatial dissimilarity of CAPs between groups. Figure 3 shows the comparison of temporal and spatial properties of matched CAPs between the WT and TG. Only the first CAP displayed a higher median occurrence in the TG as compared to WT while the duration and occurrence rates of all other CAPs didn’t show any significant difference. CAPs 2, 3, 5 and 6 showed a significantly altered lateralization of average positive activation per voxel between the groups.

**Figure 3:**
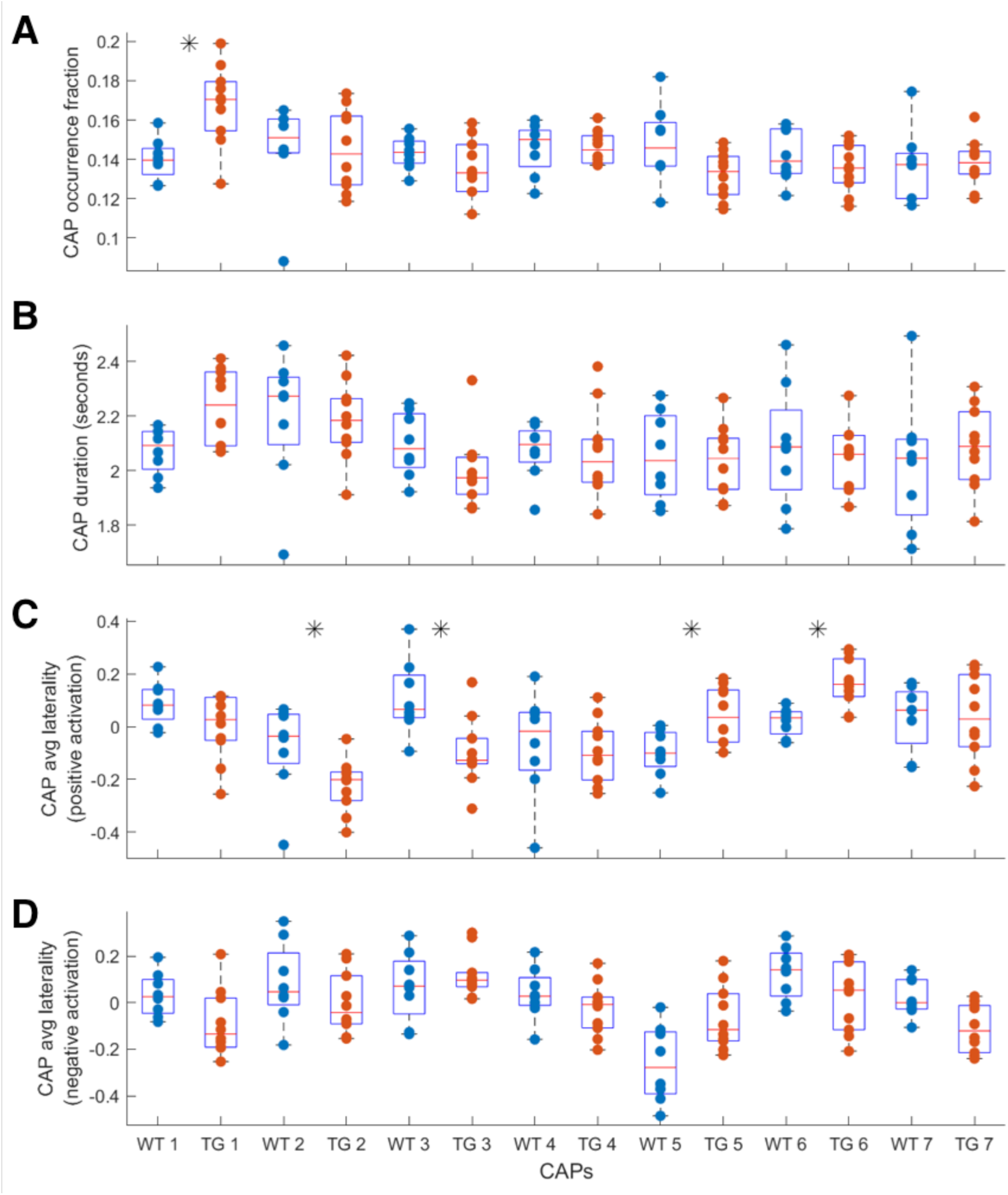
Comparison of median occurrence rate **(A)**, duration **(B)**, hemispheric lateralization of average positive activation per voxel **(C)**, and hemispheric lateralization of average negative activation per voxel **(D)** between WT and TG for each of the 7 CAPs extracted from group-level image-series. Black asterix indicates significant difference (p < 0.05, two sample rank sum test; FDR corrected for multiple comparisons with Benjamini-Hochberg correction). Most significant differences are found for the lateralization of co-activation (T > 0) in case of 4 out of 7 CAPs.

Next, we compared the properties of matched CAPs between the two groups in all partitions with number of clusters ranging between 2 and 20. Figure 4 plots the p-values of comparison of temporal (panels A and B) and spatial (panels C and D) properties. While very few CAPs, across partitions, showed significant difference in either the duration or the occurrence, 57 and 52 CAPs across multiple partitions showed significant between-group difference in the lateralization of the average positive and negative activation per voxel respectively (p < 0.05, FDR corrected for number of comparisons within each partition). We identified two prototype patterns with spatially similar representations in 7 different partitions that showed significantly higher right-hemisphere lateralization of positively activated voxels in the TG group (Figure 5). CAP 3 of 7 (Figure 2C) belonged to the first prototype pattern (Figure 5A) while CAP 2 of 7 belonged to the second group (Figure 2B, 5B). Similarly, two prototype patterns with spatially similar representations in 6 different partitions displayed significantly higher right-hemisphere lateralization of co-deactivated (T < 0) voxels in the TG animals in comparison with WT (Figure 6). The first prototype pattern (Figure 6A) was highly similar to CAP 3 of 7 (Figure 2C) while the second pattern closely matched CAP 1 of 7 showing co-deactivation of DMN-like network.

**Figure 4:**
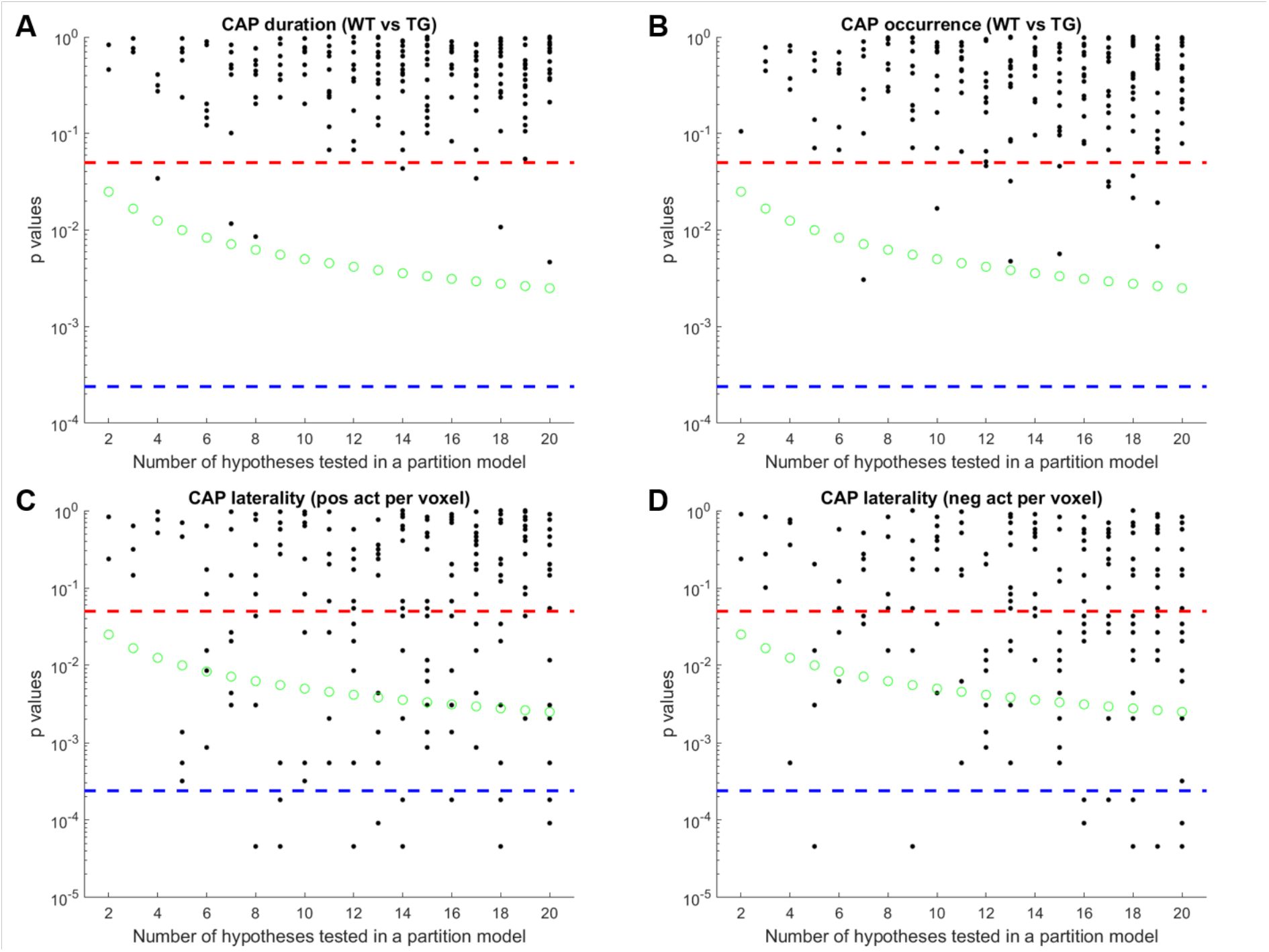
Each panel of the figure shows the p-values of comparison of mean duration **(A)**, occurrence rate **(B)**, laterality of average positive activation per voxel **(C)**, and, laterality of average negative activation per voxel **(D)** between WT and TG groups for each CAP in each partition with number of clusters ranging from 2 to 20. Thus each black marker represents a CAP. The red dashed line represents a p-value of 0.05, uncorrected. The green marker represents Bonferroni corrected threshold p-value for each partition. The blue dashed line represents the Bonferroni corrected threshold p-value across partitions. Here, CAPs are extracted from the group-level image-series.

**Figure 5:**
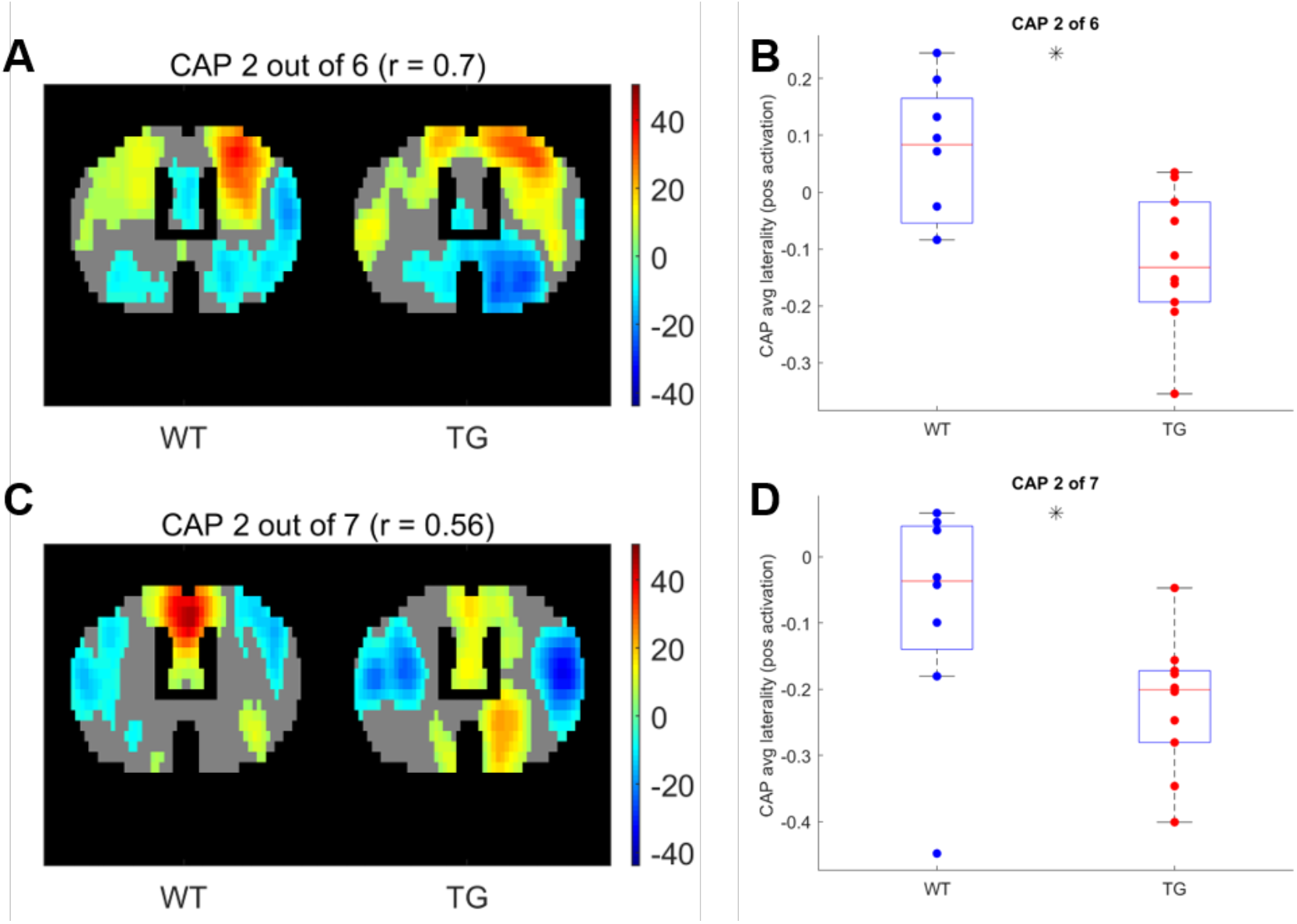
One-sample T-test map of an example CAP for two patterns (A, C respectively) with representations in 7 different partitions that show significantly higher right-lateralization (B, D respectively) of significantly (p < 1E-5; Bonferroni corrected) co activated (T > 0) voxels.

**Figure 6:**
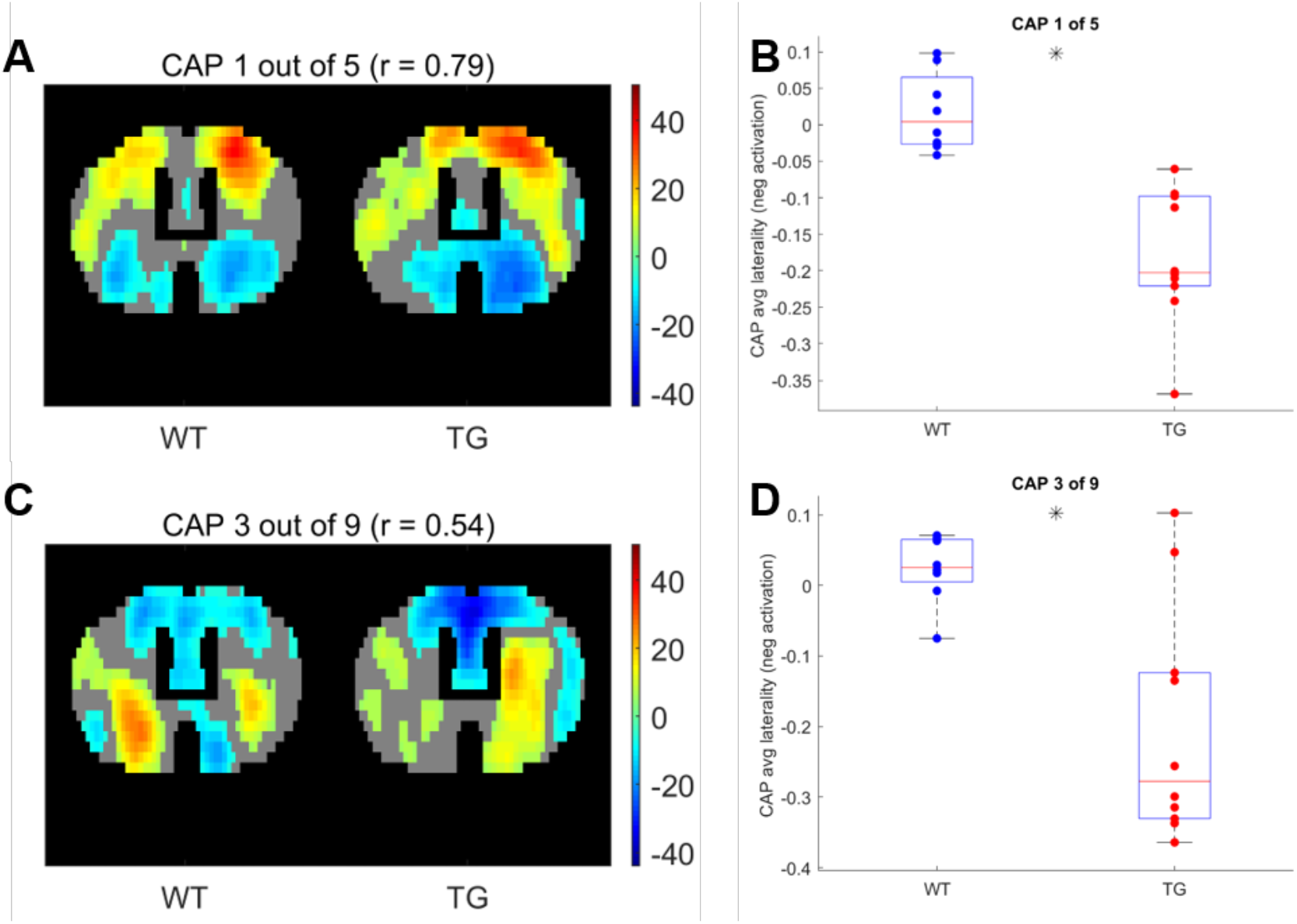
One-sample T-test map of an example CAP for two patterns (A, C respectively) with representations in 7 different partitions that show significantly higher right-lateralization (B, D respectively) of significantly (p < 1E-5; Bonferroni corrected) co deactivated (T < 0) voxels.

Finally, we tested an alternative strategy to extract CAPs. Instead of extracting CAPs separately for each group’s image-series, we concatenated both group-level image-series into a single one and applied the clustering algorithm to it. The rationale behind this data-driven approach was to identify CAPs without specification of group identity. This approach clearly didn’t require any matching as each CAP would have representations in each group. After extracting the CAPs in this manner we compared their properties between groups as before and found that both the temporal and the spatial properties of most CAPs didn’t show any significant difference (Figures 7 and S5). The failure of this approach to identify any inter-group differences can be attributed to the low statistical power in our dataset and to the fact that the clustering puts relatively more emphasis on capturing the inter-group variance than the within-group variance.

**Figure 7:**
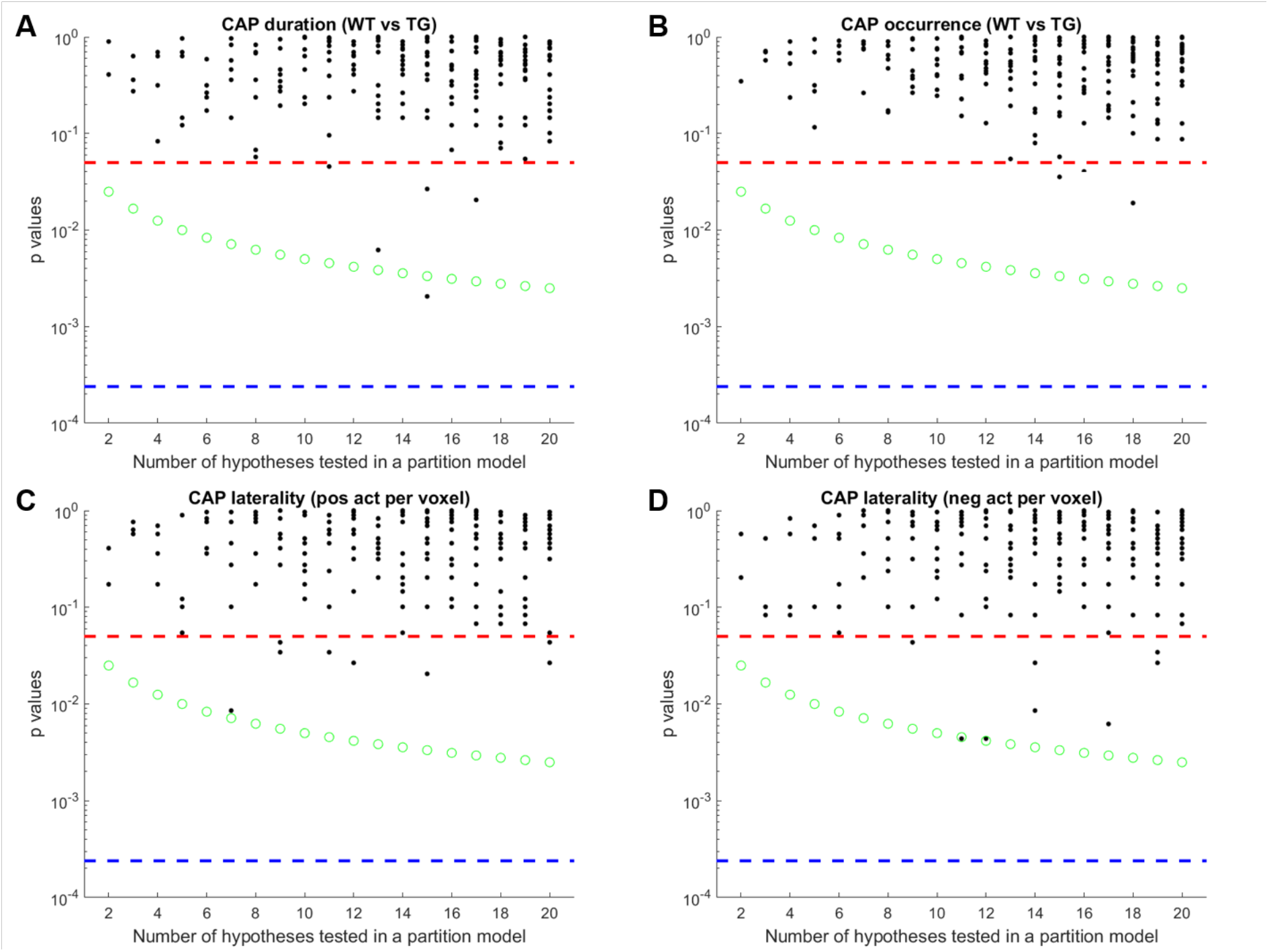
Each panel of the figure shows the p-values of comparison of mean duration **(A)**, occurrence rate **(B)**, laterality of average positive activation per voxel **(C)**, and, laterality of average negative activation per voxel **(D)** between WT and TG groups for each CAP in each partition with number of clusters ranging from 2 to 20. Thus each black marker represents a CAP. The red dashed line represents a p-value of 0.05, uncorrected. The green marker represents Bonferroni corrected threshold p-value for each partition. The blue dashed line represents the Bonferroni corrected threshold p-value across partitions. Here, CAPs are extracted from a single image-series formed by concatenating both group-level image-series.

### 3.3 Classification using CAP metrics

Finally we turned our attention to investigating the predictive power of CAP metrics to distinguish TG animals from WT. As mentioned in the methods, at first, we considered the properties of training subjects’ CAPs, extracted from the image-series of all subjects, as features to train the classifier. Figure 8A-B show the mean classification accuracy with temporal and spatial properties of CAPs respectively and their comparisons with mean chance-level accuracy as a function of partitions of the image-series. Occurrence rates and durations of strongly (r > 0.5) spatially matched CAPs performed better than the chance-level only in a few partitions. On the other hand, the spatial component performed significantly better than the chance level for all partitions with an average accuracy of 90% in all cases barring two partitions. Figure 8C-D show the confusion matrices for each of the 19 partitions. We find that the significantly greater than the chance-level accuracy observed with the temporal features of matched CAPs in 8 partitions can be attributed to primarily the WT class being better predicted. In these partitions, the prediction scores for the transgenic animals is typically lower than that for WT. On the other had, the high classification accuracy of the spatial component of matched CAPs is due to excellent predictions of both classes, albeit, it’s the TG that is predicted perfectly while some of the WT subjects are incorrectly predicted as TG. This observation could be explained by the fact that these animals are very old and hence, the some of WT animals could show patterns that are spatially very similar to the TG subjects.

**Figure 8A, B:**
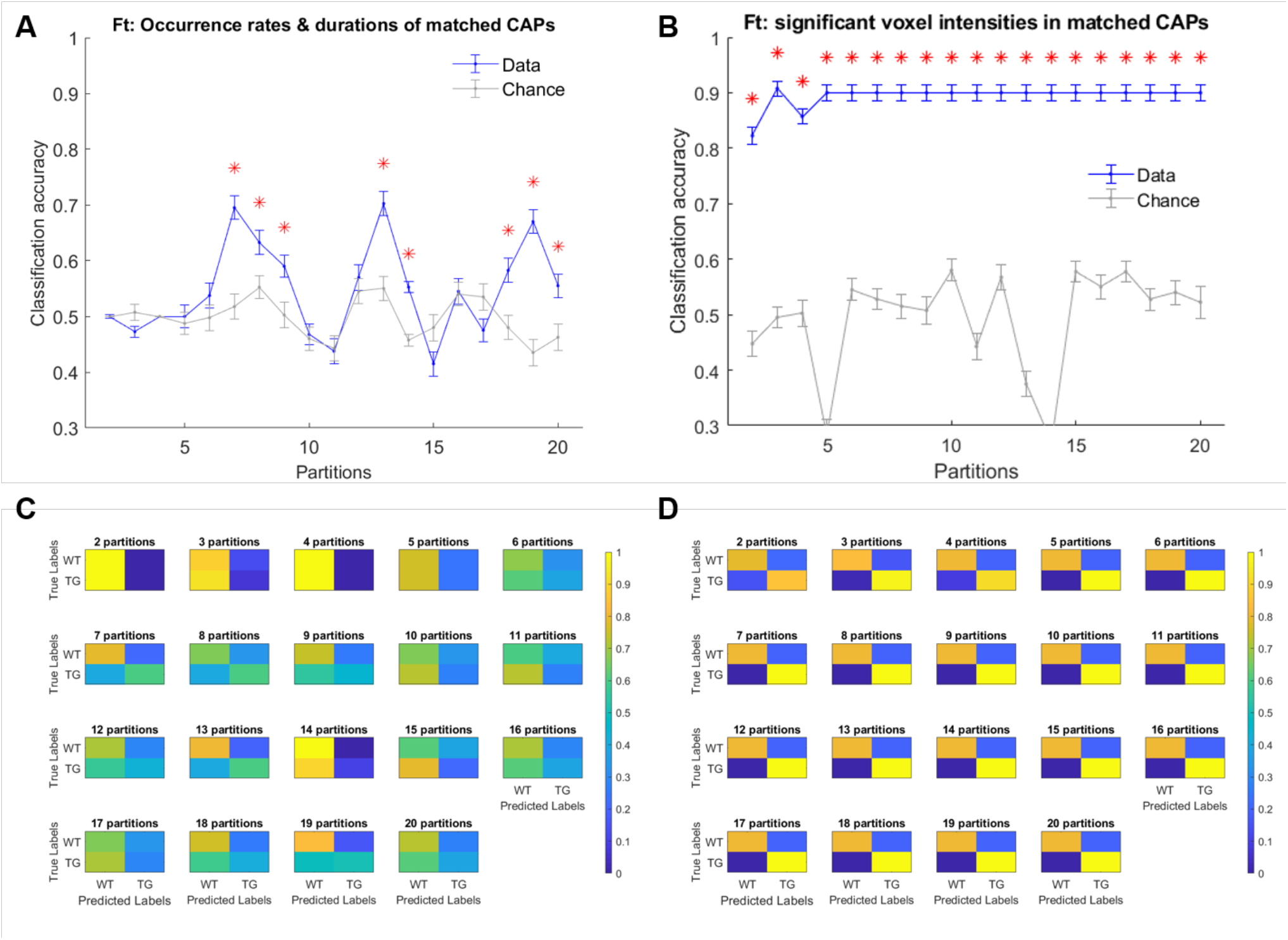
Mean accuracy of classification (blue) and mean chance-level accuracy (grey) as a function of partitions with number of clusters ranging between 2 to 20. MLR classifier is trained using duration and occurrence rates of matched CAPs (with Pearson’s correlation higher then 0.5) **(A)**, and, BOLD signal intensities of voxels with activations significantly different from zero found in the group-level WT or TG CAP **(B)**. Red asterix indicates significantly higher mean accuracy than the chance level, FDR corrected for 19 comparisons using Benjamini-Hochberg correction. **C, D:** Confusion matrices showing the scores of prediction of validation set labels of each group, pooled across 100 validations sets, for each of the 19 partitions with temporal **(C)** and spatial **(D)** aspects of CAPs.

As described in the methods section, the fact that all subjects’ image-series were used in the extraction of the CAPs could influence the classifier to predict the validation set subjects more accurately than in the case when only training set subjects’ images were concatenated to extract the CAPs. Therefore, we extracted the WT and TG group-level CAPs from the image-series of only the training set subjects and then, for every group-level CAP, obtained the WT-like and TG-like CAP for each subject (see Methods for details). Taking the spatial and temporal components of WT-like and TG-like CAPs as features we trained the classifier and tested on the validation set. As figure 9A shows, the classification accuracy with temporal features of WT-like and TG-like CAPs was close to the chance level for all partitions. Only in six partitions, the mean accuracy was significantly higher than the chance-level while never crossing a 60% mark. In contrast, the prediction accuracy of spatial features was near 100% in case of all partitions except the first two (Figure 9B).

**Figure 9A, B:**
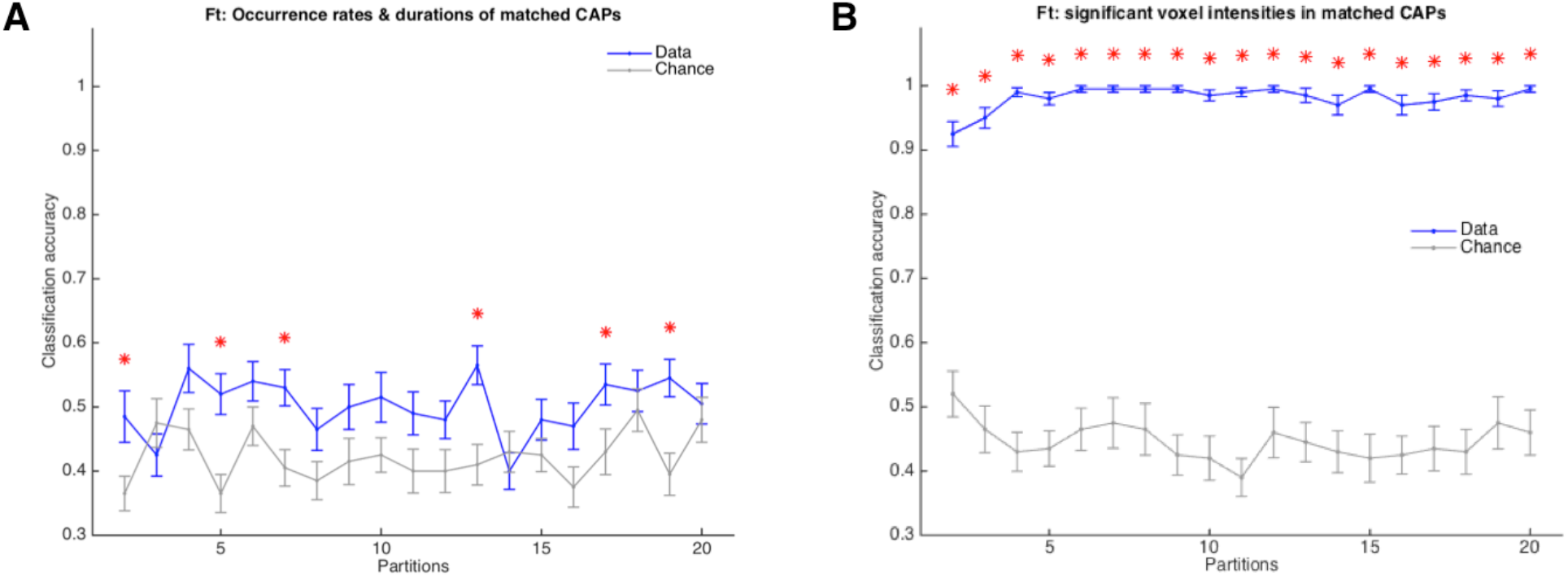
Mean accuracy of classification (blue) and mean chance-level accuracy (grey) as a function of partitions with number of clusters ranging from 2 to 20. MLR classifier is trained using duration and occurrence rates of WT-like and TG-like CAPs **(A)**, and, their BOLD signal intensities of voxels with activations significantly different from zero found in the group-level WT or TG CAP **(B)**. Red asterix indicates significantly higher mean accuracy than the chance level, FDR corrected for 19 comparisons using Benjamini-Hochberg correction. Here the group-level WT and TG CAPs are extracted from group-level image-series formed from only the training set of subjects.

## 4 Discussion

In this article, we investigated spatial and temporal properties of resting-state co-activation patterns extracted in a mouse model of Alzheimer’s disease at a very old age. We found very few inter-group differences in the temporal component of CAPs. More robust differences were found in the hemispheric lateralization of co-activations and co-deactivations of brain regions in multiple CAPs. Typically, both the co-activations and co-deactivations were significantly more right lateralized in the diseased animals as compared to the healthy. These differences, especially in the spatial features of CAPs, suggested that they could serve as accurate predictors of the disease. We therefore tested the predictive ability of both the spatial and temporal features of CAPs using a supervised learning approach. The prediction accuracy of temporal features was near the chance level while the spatial features distinguished the diseased animals from healthy control with near perfection.

### Methodological considerations

CAPs were obtained using a recently developed approach (Gutierrez-Barragan et al. 2019) in which all time-frames, as opposed to only those corresponding to the supra-threshold BOLD signal in a seed region, are clustered in each group separately, followed by identifying the optimal number of clusters at which the across-subject variance saturates. In our case, the identification of elbow point in the plot of explained variance versus the number of clusters was sub-optimal as the variance increased monotonically with the number of clusters and the fraction of variance gain didn’t drop significantly after a specific partition as was the case in the article by Gutierrez-Barragan et al. This could be due to lower statistical power in our cohort (8 WT and 10 TG animals). We therefore continued identification of CAPs for all partitions with number of clusters ranging between 2 and 20 and compared the properties of only those CAPs that showed significantly higher spatial similarity between the WT and the TG group than that arising by chance.

We also extracted CAPs in the same range of partitions from a combined image-series formed by concatenating images from animals of both groups. Here, we did not find significant differences in either the temporal or spatial features of CAPs. This finding could be attributed mainly to the nature of the clustering algorithm; K-means appears to identify clusters that are similar across both groups thereby putting more emphasis on explaining across group variance than within group. Low statistical power in each group could also explain this observation as insufficient variability within group would mean the clustering would fail to find a group-specific pattern as a separate cluster and combine it instead with a larger cluster of similar observations across groups. It would be interesting to test, in a cohort of sufficiently large datasets, if both group-level and combined approaches yield similar results.

In order to do the classification we used, as a classifier, multinomial logistic regression with regularization. MLR has been shown to be efficient classifier for categorical variables as was the case in this study (Pallarés et al. 2018). At first, we performed the classification by using just the representations of group-level CAPs, obtained by analyzing data from all subjects, in each individual subject. However, in order to mimic a more clinical setting in which only the training data would be available to identify group-level CAPs and the purpose of the marker would be to diagnose a test participant, we used only the training set animals’ data to extract the group-level WT and TG CAPs. Subsequently we obtained the subject-level CAPs in training as well as the test-set subjects by seeding the clustering algorithm with initial centroids formed by averaging over local peaks of correlation with the corresponding group-level CAPs.

### CAPs Topology

Out of the seven CAPs, we found two pairs of spatially anti-correlated CAPs. The first pair showed co-activation and co-deactivation of cingulate and motor cortices that were anti-correlated with activations and deactivations of the striatum and somatosensory cortices. Both these pairs of CAPs were similar to the first four CAPs obtained by Gutierrez-Barragan et al. (2019) although the extent of significantly co-activated voxels was much less. This observation could be attributed to the lower statistical power in our cohort. Cingulate cortex is a prominent region in the mouse DMN-like network while somatosensory cortices belong to the LCN (Gozzi and Schwarz 2016). The explanation for co-activation of motor cortices with DMN-like network, also found in the QPPs extracted in this cohort (Belloy et al. 2018), could lie in the age of the animals as human studies have shown that resting-state network segregation is reduced with ageing (Chan et al. 2014; Vidal-Piñeiro et al. 2014).

### Resting-state markers of Alzheimer’s disease

Human studies of the resting-state in patients with Alzheimer’s disease have mostly focused on static functional connectivity (FC) analyses (Badhwar et al. 2017). Altered DMN FC has been the most consistent finding in these studies (Mevel et al. 2011). Since regions constituting the DMN (posterior cingulate cortex, in particular) are also the targets of AD in terms of deposits of amyloid-beta plaques, alterations in RSN-FC have been shown to correlate with these deposits especially in patients with high amyloid burden(Sperling et al. 2009; Koch et al. 2015; Myers et al. 2014). These changes in the DMN-FC of animal cohort used in our study, were observed and confirmed in the previous work by Belloy et al.(2018). Going beyond the static FC, Belloy et al. showed that short (3s), spatio-temporal patterns of recurring neural activity called the quasi-periodic patterns (QPPs) contributed to the FC changes. They also found that group-specific QPPs occurred less frequently in the other group’s image-series and that the dominant QPPs from each group were anti-correlated to each other. A recent study (Ma et al. 2020) investigated co-activation patterns in healthy elderly participants, patients with mild cognitive injury (MCI) and AD patients and found that average dwell time in the DMN was reduced in AD. In contrast to both these studies, we found significant differences in the spatial, rather than temporal, component of CAPs in terms of hemispheric lateralization of co-activation and co-deactivation. We found significantly higher right-lateralization in the TG animals for both co-activated as well as co-deactivated voxels. In humans, several restingstate networks show lateralizations that depends on age and gender (Agcaoglu et al. 2015). In fact, significant reduction in the lateralization of the DMN has recently been reported for groups of amyloid beta positive patients of MCI or dementia when compared with a group of amyloid beta negative participants with no cognitive impairment (Banks et al. 2018). In mice, a strong bilateral organization of resting-state networks, found using independent component analysis, has been reported (Grandjean et al. 2020). Our finding in WT mice that shows mostly bilateral CAPs is in line with this observation. On the other hand, in mouse models of autism spectrum disorder, brain lateralization, especially in the striatum, has been reported using MRI and immunohistochemistry (Grabrucker et al. 2018). Therefore, right-lateralization of prominent CAPs in TG mice is an interesting finding that needs to be further investigated.

### Potential of CAPs as a biomarker

Spatial features of CAPs, although similar at a group-level, were sufficiently different at the subject-level as evidenced by inter-group differences in the hemispheric lateralization of activations. These differences were robust enough to make CAPs a highly accurate predictor of the disease at this late manifest stage. As mentioned before, we used two strategies with the latter being more appropriate for a clinical setting in which the classifier could be trained using CAPs extracted out of only the training dataset with the aim of diagnosing new “un-seen” participants. We found that the spatial features of CAPs predicted the identity of test-set subjects perfectly with above 99% accuracy. This result suggests that CAPs could also predict more complex scenarios such as behavioral deficits of patients at different stages or outcome of treatments thereby making resting-state CAPs a very promising candidate for a biomarker for Alzheimer’s disease as well as other neurodegenerative diseases.

## 5 Author Contributions

M.H.A. designed research, designed analysis tools, performed research, analyzed data and wrote the manuscript. M.E.B. acquired the RS-fMRI data and provided inputs on the manuscript. A.V.d.L., G.A.K. and M.V. designed research and provided inputs on data analysis and the manuscript.

